# Exploring the microbiome-wide lysine acetylation, succinylation and propionylation in human gut microbiota

**DOI:** 10.1101/2020.12.14.422746

**Authors:** Xu Zhang, Kai Cheng, Zhibin Ning, Janice Mayne, Krystal Walker, Hao Chi, Charles L. Farnsworth, Kimberly Lee, Daniel Figeys

## Abstract

**Background:** Lysine acylations are important post-translational modifications that are present in both eukaryotes and prokaryotes, regulating diverse cellular functions. Our knowledge of the microbiome lysine acylation remains limited due to the lack of efficient analytical and bioinformatics methods for complex microbial communities.

**Results:** We show that serial enrichment using motif antibodies successfully captures peptides containing lysine acetylation, propionylation and succinylation from human gut microbiome samples. A new bioinformatic workflow consisting of unrestricted database search confidently identified >60,000 acetylated, and ~20,000 propionylated and succinylated gut microbial peptides. Characterization of these identified modification-specific metaproteomes, i.e. meta-PTMomes, demonstrates that lysine acylations are differentially distributed in microbial species with different metabolic capabilities.

**Conclusion:** This study provides an analytical framework, consisting of a serial immunoaffinity enrichment and an open database search strategy, for the study of lysine acylations in microbiome, which enables functional microbiome studies at the post-translational level.

## Background

The human microbiome, an assembly of microorganisms that reside on the surface of our body, is increasingly recognized as a prominent factor affecting human health and the development of diseases, such as inflammatory bowel diseases, metabolic disorders, cardiovascular diseases, and neurological disorders. Accumulating evidence also suggests that the human microbiome, in particular the gut microbiome, determines the toxicity/efficacy of many drugs, such as digoxin and L-dopa [1-3], and affects patient responses to therapeutics, such as PD1 blockade anticancer therapy [4, 5]. These striking findings make the microbiome a non-negligible aspect in current biomedical research. However, our knowledge of the microbiome remains limited, and in particular on the diverse metabolic pathways in our gut microbiome, how they are regulated, and whether the regulation processes change in diseases. For example, only a few of the many previously reported drug-microbiome interactions have well characterized mechanisms and drug metabolism pathways [6, 7]. More functional approaches such as metatranscriptomics, metaproteomics and metabolomics, have been used to study the human microbiome [8]. These meta-omics approaches can provide a better understanding of the functional or metabolic alterations of the microbiome in disease or in response to drug treatment. However, how the microbiome metabolic pathways are regulated remains tremendously underinvestigated.

Post-translational modification (PTM) is an important molecular regulatory mechanism in all kingdoms of organisms, and a few PTMs, such as acetylation and methylation, have been reported to be abundant in human and environmental microbiomes [9-11]. We have previously reported that the integration of immunoaffinity enrichment and mass spectrometry allowed the identification of >35,000 lysine acetylated (Kac) peptide in gut microbiome [9]. The application of this lysine acetylomic approach for the study of microbiomes in pediatric patients with Crohn’s disease (CD) demonstrated a down-regulation of lysine acetylation levels in metabolic pathways for the production of short-chain fatty acids (SCFAs) in CD compared to control [9]. Moreover, lysine acetylation occurs on all key enzymes in SCFA metabolism pathways of the microbiome, including the enzymes for generating succinate, propionate or their derived metabolites, such as succinyl-CoA and propionyl-CoA [9]. These findings suggest that the enzymes for gut microbial SCFA metabolism can be regulated by acetylation, succinylation and propionylation, three types of lysine acylations that are reported to be frequently present on bacterial proteins and may interact with each other in regulating cellular metabolisms [12, 13].

SCFAs are energy sources for intestinal epithelium cells and known to protect intestinal barrier function, and different SCFAs (i.e., acetate, butyrate or propionate) may have different activities [14]. The findings from the above microbiome-wide lysine acetylomic study highlight the importance of lysine acylations in regulating the microbial functions, and the study of the crosstalk between these acylations may provide insights on how the metabolic outcomes of SCFA metabolism are regulated in microbiome. Unfortunately, there is no published study on the different lysine acylations in microbiome to date nor established technique enabling efficient microbiome-wide study of multiple PTMs. Therefore, in this study we developed an analytical workflow by integrating serial immunoaffinity enrichment with high resolution mass spectrometry and unrestricted metaproteomic database search strategy, which enables efficient and comprehensive identification and quantification of lysine acetylation, propinoylation and succinylation in microbiomes. This analytical workflow provides a framework for high throughput and deep –omic study of PTMs in the microbiome (termed meta-PTMomics), complementing the current toolbox for functional microbiome study.

## Results and Discussion

### Open search enables the identification of both expected and unexpected modifications in meta-PTMomics

An important challenge for meta-PTMomics studies is controlling the false discovery rate when using a tremendously expanded database search space for modified peptide identification (i.e., large database and more variable modifications). In our previous study, we adopted an iterative database search strategy to generate a refined smaller database used for the identification of lysine acetylated peptides with a restricted search strategy, i.e. Andromeda [15]. More recently, open search strategies for mass spectrometry data were reported for the identification of both expected and unexpected protein modifications, with increased sensitivity, accuracy and numbers of peptide identification while controlling false discovery rates [16]. Several ultra-fast open search engines, such as Open-pFind and MSFragger [17, 18] were reported, which greatly increased the applicability of open search in proteomics studies, including metaproteomics [10].

To evaluate the performance of open search for meta-PTMomics analysis, we reanalyzed our previous Kac dataset using an open search workflow implemented in the MetaLab software tool (www.imetalab.ca) (Fig. 1A; details in Methods section). Briefly, we first clustered the spectra from all samples, including Kac aliquots and metaproteomic aliquots. Then the clustered spectra were searched against the human gut microbial IGC (integrated gene catalog) database using Open-pFind to generate a refined database [18]. The refined database was then used for peptide identification of each sample using Open-pFind and quantification using FlashLFQ [19]. Using this open search approach, we identified 1.6 times more Kac peptides than our previous study, yielding a total of 57,406 Kac peptides identified from six microbiome samples. Moreover, in addition to Kac peptides, we also identified other lysine acylated peptides, including 1,263 lysine propionylated (Kprop) peptides, 338 lysine formylated (Kform) peptides and 79 lysine butyrylated (Kbut) peptides. These lysine acylated peptides are likely co-enriched during Kac immunoaffinity enrichment. Along with other modifications (e.g., oxidation, deamidation, Gln->pyro-Glu, carboxymethylation, and carbamylation) that were also identified in unenriched samples (Table S1), an average spectra identification rate of 43% and 57% was achieved in Kac and metaproteomic datasets, respectively. Altogether, these results suggest that the open search strategy is well-suited for enrichment-based microbiome-wide PTM analyses, enabling the identification of both expected and unexpected modifications.

**Figure 1.**
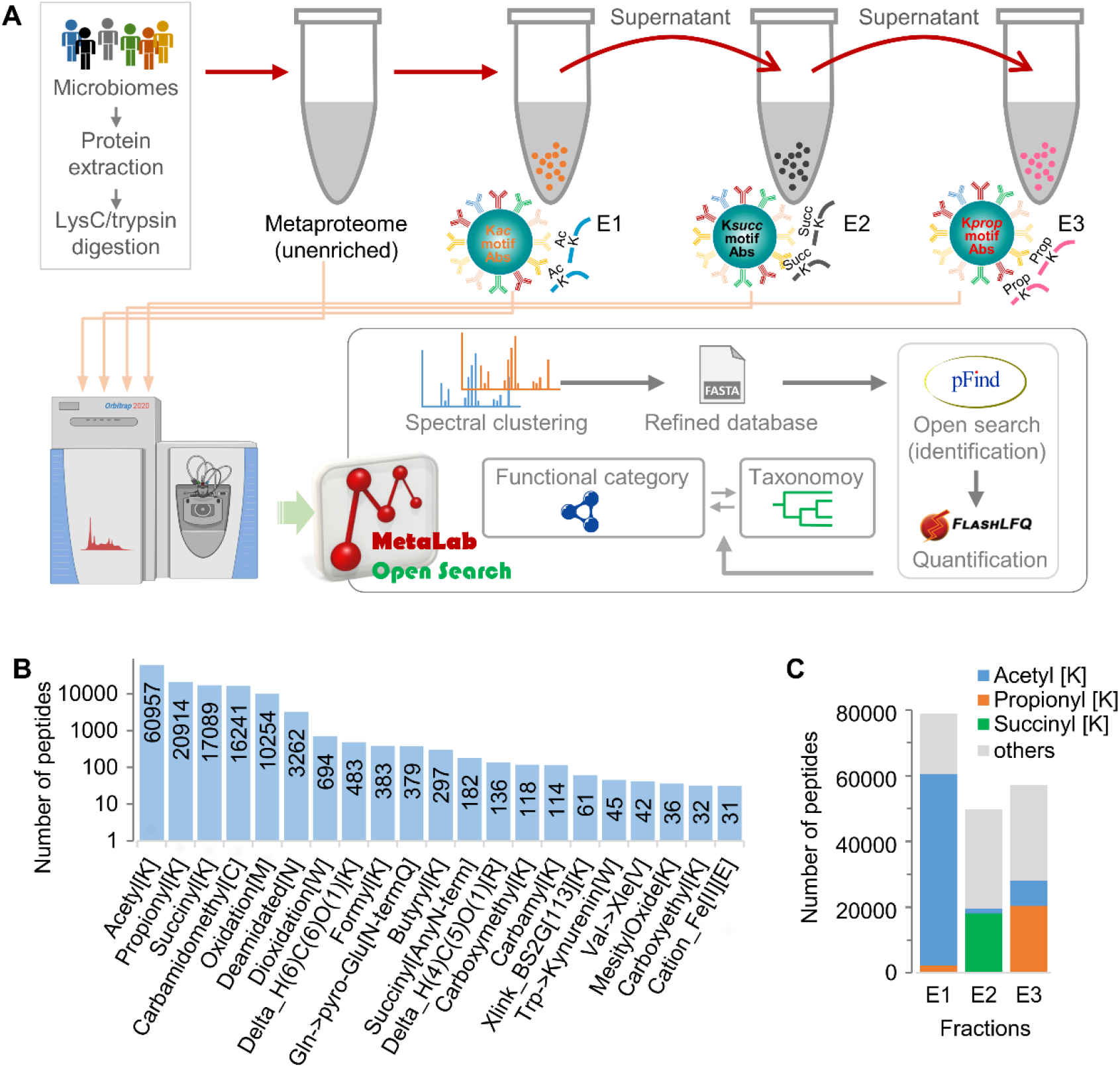
Serial immunoaffinity enrichment and open search approach for meta-PTMomic study of lysine acylations. (A) Experimental and bioinformatics workflow. Fecal microbiomes were collected from adult volunteers and their proteins were extracted and digested using lysC and trypsin. The resulting peptides were then used for a serial enrichment using Kac, Ksucc and Kprop motif antibodies, respectively, and the eluted peptides were measured using mass spectrometer (MS). MS data was analyzed using MetaLab open search workflow, consisting of (1) spectral clustering and database search to generate a refined database, (2) database search using Open-pFind, (3) quantification using FlashLFQ, and (4) taxonomic and functional annotation for all identified proteins/peptides. (B) Distribution of identified modifications. Modifications with >30 modified peptides identified were shown. (C) Distribution of identified peptides in each enrichment fractions. Unmodified peptides and other less frequently identified modifications were combined as other in the plot.

### A serial enrichment-based framework for deep meta-PTMomics study in microbiomes

To concurrently investigate multiple lysine acylations in microbiomes, we applied serial immunoaffinity enrichments of different lysine acylation-modified peptides from the same sample [20]. Briefly, in this workflow, the proteins from the gut microbiome are extracted using urea-free lysis buffer, digested using trypsin/lysC, and the resulting peptides are subjected to a serial enrichment for lysine acetylation, succinylation and propionylation using PTMscan® motif antibody kits (Cell Signaling Technology, Inc) (Fig. 1A). The eluted enriched peptides are analyzed using mass spectrometry and modified peptides/proteins are identified using the open search strategy described above. Overall, across six human microbiomes, a total of 60,957 Kac, 20,914 Kprop and 17,089 Ksucc peptides were identified, along with other lysine acylated peptides (i.e., 383 Kform and 297 Kbut peptides). Kac, Kprop and Ksucc are also the three most frequently identified modifications in this enrichment dataset (Fig. 1B), suggesting successful enrichment of lysine acylated peptides using motif antibodies. Interestingly, 2,284 Kprop peptides were identified in the Kac-enriched fraction (E1), representing 3% of all identified modified peptides in this fraction. Similarly, 1,552 Kac peptides were also identified in the Ksucc-enriched fraction (E2, 3%), and 7,546 Kac peptides were identified in Kprop-enriched fraction (E3, 13%) (Fig. 1C). These findings suggest a high degree of co-enrichment using the immunoaffinity enrichment approach, in particular for the modifications with similar size, i.e., Kac and Kprop.

The serial enrichment approach adopted in this study allows direct comparison of the abundance of each modification in the same sample. We therefore estimated the abundance of each modification using the total modified peptide intensity, which showed that Kac peptides were the most abundant lysine acylation with around 5-fold higher intensity than Ksucc or Kprop peptides (Fig. S1). Motif analysis using pLogo showed marked motif differences between the three modifications (Fig. S2). In agreement with our previous observation [9], Kac motifs were featured with E and D being the most frequent amino acids at −1 position. For Kprop sites, glycine and proline occurred most frequently at the +1 position, and alanine occurred frequently at −2 position. Alanine was also the most frequent amino acid surrounding Ksucc sites at most positions within the observation window. The amino acid preferences for each modification didn’t likely result from the bias of antibodies used for enrichment because different motifs were observed for mouse liver samples using the same kits (results not shown; from Cell Signaling Technology, Inc).

### Widespread co-modification of lysine acylations in gut microbial metabolic pathways

To examine the extent of co-modification or competitive modification between the three lysine acylations, we first directly compared the peptide sequences of Kac, Kprop and Ksucc peptides identified in this study. Results showed that >50% of the identified Kprop and Ksucc peptide sequences can also be modified by acetylation and 4,428 (6.5%) peptide sequences can be modified by all three acylations (Fig. 2A). In agreement with peptide sequence analysis, the overlap at protein and modification site levels showed that acetylation happens at 88% and 89% of the identified Kprop and Ksucc proteins, and 66% and 72% of the Kprop and Ksucc sites, respectively (Fig. S3). Comparison of the top 100 abundant modified peptides of the three modifications showed higher overlap between Kac and Kprop, and the least overlap between Kac and Ksucc (Fig. 2B). Ten peptide sequences were among the top 100 abundant peptides of all three modifications, including 3 peptides from the protein phosphoenolpyruvate carboxykinase (PCK) as well as peptides from other enzymes both upstream (i.e., phosphoglycerate kinase, 3-phosphoglycerate dehydrogenase, phosphofructokinase, and glyceraldehyde-3-phosphate dehydrogenase) and downstream (i.e., methylmalonyl-CoA mutase) of the SCFA metabolism pathway (Fig. 2B). These results further show that the glycolysis and SCFA metabolism are among the most abundantly acylated metabolic pathways in the gut microbiomes.

**Figure 2.**
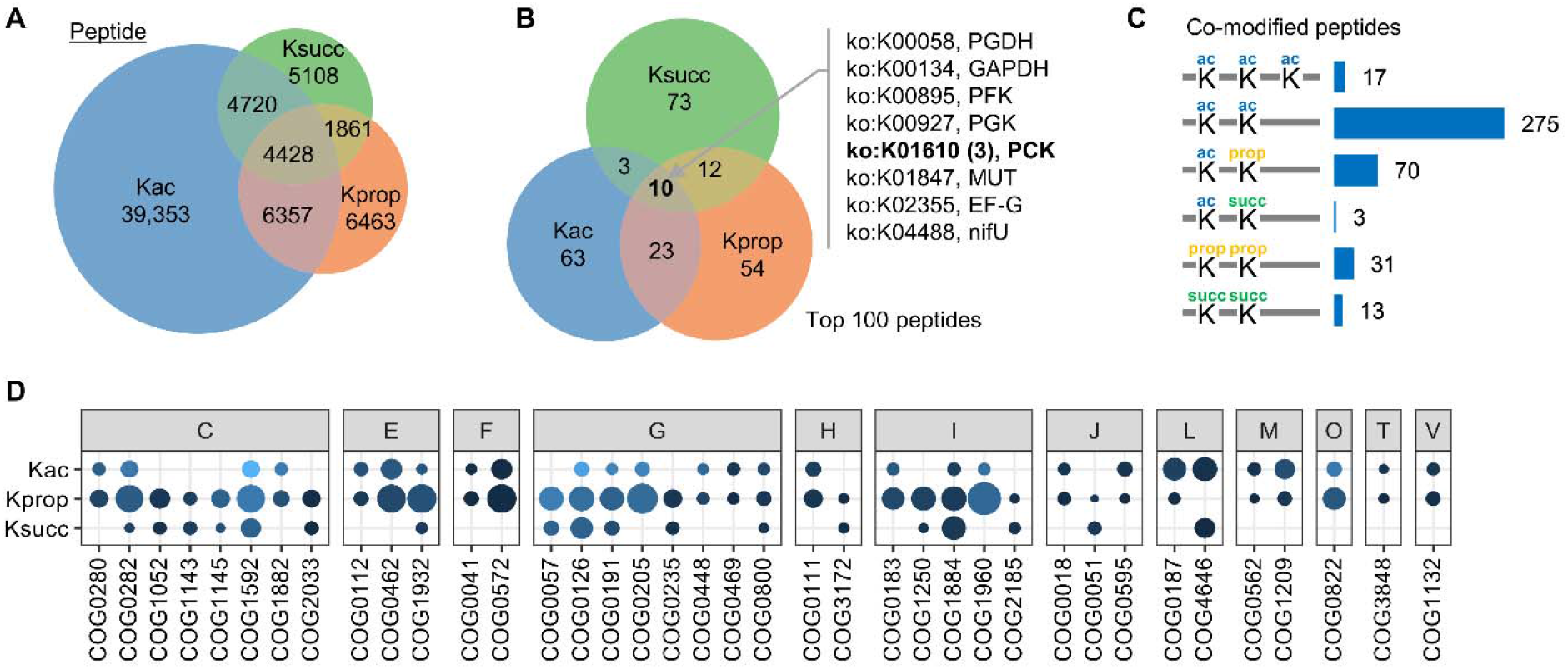
Functional characterization of identified lysine acylated peptides and proteins. (A) Overlap of the identified Kac, Kprop and Ksucc peptide sequences. (B) Overlap of the top 100 abundant Kac, Kprop and Ksucc peptide sequences. The KEGG Orthology IDs and names of the ten peptide sequences that overlap among all three modifications were shown. PGDH, 3-Phosphoglycerate dehydrogenase; GAPDH, glyceraldehyde-3-phosphate dehydrogenase; PFK, phosphofructokinase; PGK, Phosphoglycerate kinase; PCK, phosphoenolpyruvate carboxykinase; MUT, methylmalonyl-CoA mutase; EF-G, elongation factor G; nifU, nitrogen fixation protein NifU. (C) Number of identified peptides with two or more lysine acylation modifications. (D) Significantly enriched COGs by Kac, Kprop or Ksucc proteins identified in microbiome. COGs were organized according to the COG category: C, Energy production and conversion; E, Amino acid metabolis and transport; F, Nucleotide metabolism and transport; G, Carbohydrate metabolism and transport; H, Coenzyme transport and metabolism; I, Lipid transport and metabolism; J, Tranlsation; L, Replication and repair; M, Cell wall/membrane/envelop biogenesis; O, Post-translational modification, protein turnover, chaperone functions; T, Signal transduction mechanism; V, Defense mechanisms. The size of the dot indicates the –log10(P value) and color indicates the number of hits (low, light blue; high, dark blue).

Among the identified lysine acylated peptides in microbiomes, 275 peptides had two Kac sites, 17 had three Kac sites (Fig. 2C). In addition, we also identified 70 peptides that were co-modified by Kac and Kprop, 3 peptides that were co-modified by Kac and Ksucc (Fig. 2C). These findings suggest potential cross-talk between these lysine acylations in regulating the functions of proteins. Interestingly, the 70 Kac/Kprop co-modified peptides were mainly distributed in Firmicutes species, such as *Faecalibacterium*, and enzymes with functions related to glycolysis and SCFA metabolism, such as phosphoglycerate kinase (PGK), 3-hydroxybutyryl-CoA dehydrogenase (HBD) and phosphoenolpyruvate carboxykinase (PCK). To further explore the functional distribution of the three modification-specific metaproteomes (termed meta-PTMomes), we performed functional enrichment analyses for all identified Kac, Kprop and Ksucc proteins, which showed that functional categories X (Mobilome), L (Replication, recombination and repair), I (Lipid transport and metabolism) and C (Energy production and conversion) were significantly enriched in all three PTM-omes, while category G (Carbohydrate transport and metabolism) was only enriched by Kprop proteins and V (Defense mechanisms) was only enriched by Kac proteins (Fig. S4). There are 127 COGs that were significantly enriched in at least one modification, 38 COGs enriched in at least two modifications, and 7 COGs enriched in all three modifications (Fig. 2D and Table S2). Interestingly, all the seven overlapped enriched COGs, namely acetate kinase (COG0282), rubrerythrin (COG1592), phosphoserine aminotransferase (COG1932), PGK (COG0126), fructose/tagatose bisphosphate aldolase (COG0191), 2-keto-3-deoxy-6-phosphogluconate aldolase (COG0800) and methylmalonyl-CoA mutase (COG1884), are functions related to key enzymes in glycolysis and the production of SCFAs, which is in agreement with the above observations on abundant peptides.

### Phylotype-specific distribution of different lysine acylations in microbiome

Taxonomic analyses of all the identified Kac, Kprop or Ksucc peptides showed that most of the lysine acylated peptides are derived from Firmicutes and Bacteroidetes in microbiome. However, Ksucc peptides were more frequently distributed in Bacteroidetes compared to Kac and Kprop peptides (Fig. 3A). The Firmicutes-to-Bacteroidetes (F/B) ratios in Ksucc-specific meta-PTMome were significantly lower than those in Kac- and Kprop-specific meta-PTMomes (Fig. 3B). Genus-level composition analyses showed that *Bacteroides* and *Prevotella*, two genera of Bacteroidetes, were the dominant genera in Ksucc-specific meta-PTMome (Fig. 3C). The relative abundance of *Faeccalibacterium*, a Firmicutes genus, was markedly lower in Ksucc-specific meta-PTMome than those in Kprop- and Kac-specific meta-PTMomes (Fig. 3C). Similar to lysine acetylation, it has been shown that bacterial protein lysine succinylation can happen non-enzymatically in the presence of succinate metabolites, such as succinyl-CoA and succinyl-phosphate [12]. This non-enzymatic protein lysine succinylation mechanism indicates that the bacterial species that can produce succinate may also have higher Ksucc levels. Accordingly, succinate producers in human gut are mainly distributed in the phylum Bacteroidetes [21].

**Figure 3.**
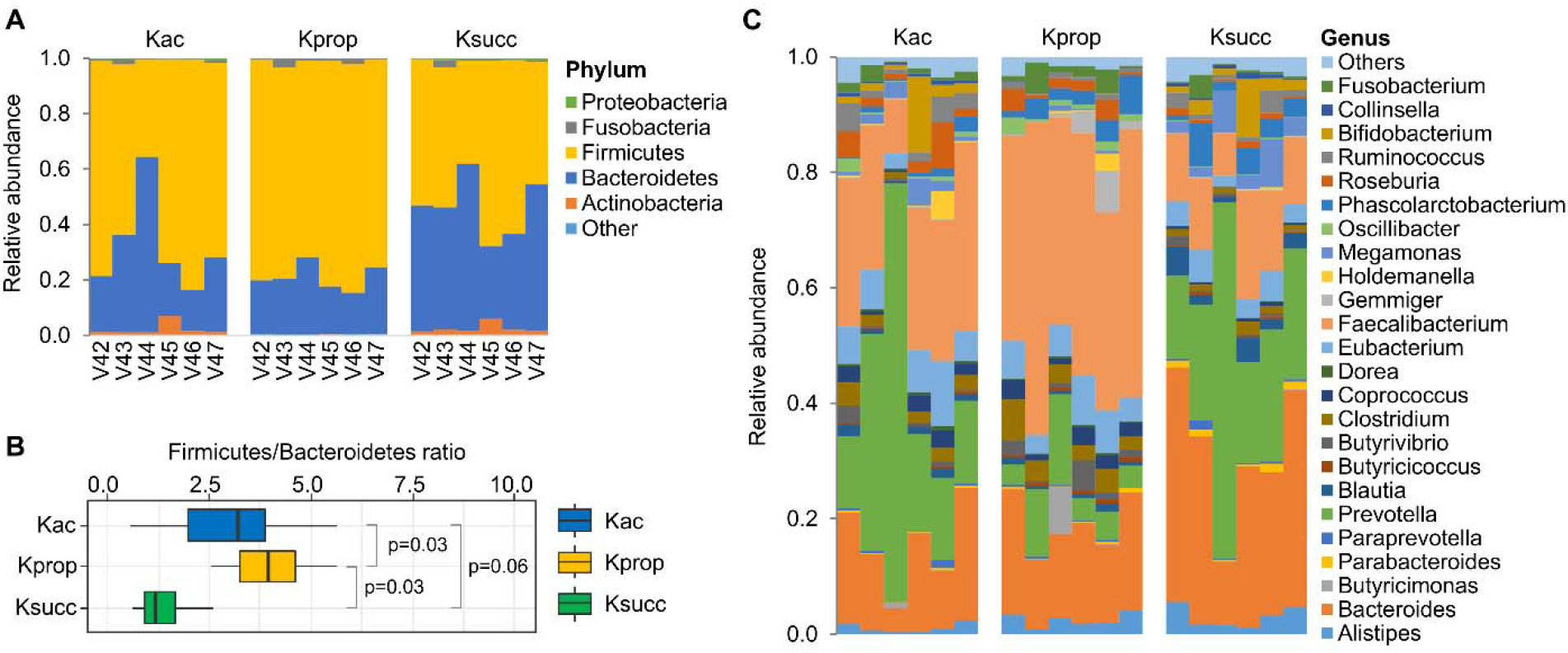
Taxonomic distribution of identified lysine acylated peptides in gut microbiome. (A) Phylum level relative abundance of quantified lysine acylated peptides in each sample. (B) Firmicutes-to-Bacteroidetes ratio of quantified lysine acylated peptides for each sample. Significance was evaluated using Wilcoxon matched-pairs signed rank test and P values were indicated. (C) Genus level relative abundance of quantified lysine acylated peptides in each sample.

These findings are in agreement with the observation of this study that identified Ksucc peptides were mainly derived from Bacteroidetes species, indicating a phylotype-specific distribution of different lysine acylations in the human gut microbiome.

## Conclusions

In summary, we showed that an integration of serial immunoaffinity enrichments, high resolution tandem mass spectrometry and unrestricted database search strategy is well suited for the comprehensive study of microbiome-wide lysine acylations. This study demonstrates that lysine acylations are widely present and may cross-talk with each other in the human gut microbiome, in particular in the metabolic pathways of glycolysis and for the production of SCFAs. In addition, the levels of different lysine acylations vary among different microbial species that encode different metabolic capabilities. Further study of the abundance alterations and/or cross-talk of these lysine acylation-specific meta-PTMomes in healthy and diseased individuals may provide more insights on how the gut microbial metabolisms are regulated, which may eventually help understand the mechanisms underlying the complex host-microbiome interactions.

## Methods

### Sample collection and processing

Fresh stool samples were collected from six healthy volunteers (age 23-28; residents at Ottawa for at least one year) with a protocol (Protocol # 20160585-01H) approval by the Ottawa Health Science Network Research Ethics Board at the Ottawa Hospital, Ottawa, Ontario, Canada. Informed consent was obtained from all volunteers. Samples were immediately put on ice after defecation and transferred to lab for processing. Briefly, around 3 g of stool was resuspended and homogenized in 30 ml cold PBS (pH 7.6) with the help of glass beads and vortexing. Debris was then removed by low speed centrifugation at 300 *g*, 5 min at 4 °C for three times. The resulting supernatant was then centrifuged at 14000 *g*, 20 min at 4 °C for harvesting microbial cells.

The microbial cells were then subjected to protein extraction using ultrasonication and lysis buffer containing 4% (w/v) SDS, 50 mM Tris-HCl (pH 8.0) and protease inhibitor (one tablet of Roche cOmplete™ mini per 10 ml lysis buffer) [22]. Ultrasonication was performed using a Qsonica Q700 (Qsonica, LLC) equipped with a cup horn for 10 min at 8 °C (amplitude, 50%; 10s pulse on and 10s pulse off). Remaining cell debris was removed by centrifugation at 16,000 *g* for 10 min. The supernatant was then precipitated by adding five volumes of ice-cold precipitation buffer (acetone/ethanol/acetic acid, 50%/50%/0.1% (v/v/v)) overnight at −20 °C. After three washes using ice-cold acetone, proteins were re-suspended in 6 M urea, 100 mM ammonium bicarbonate buffer (pH 8.0) for protein assay and proteolytic digestion.

### Proteolytic digestion and desalting

Protein concentration was determined using a DC protein assay (Bio-Rad Laboratories, Inc) according to the manufacturer’s instruction. Approximately 10~15 mg of proteins were then used for a sequential digestion using lysyl endopeptidase (Lys-C; Wako Pure Chemical Corp., Osaka, Japan) and trypsin (Worthington Biochemical Corp., Lakewood, NJ, USA). Briefly, proteins were first reduced with 10 mM dithiothreitol (DTT) and alkylated with 20 mM iodoacetamide (IAA) at room temperature; then after dilution the digestion was performed using 1 mAU of Lys-C per 50 μg protein for 4 hours followed by the addition of 1 μg of trypsin per 50 μg protein for overnight at room temperature with agitation. The resulting digests were then desalted using Waters Sep-Pak® Vac 3cc (200 mg) tC18 cartridges and eluted using 80% (v/v) acetonitrile/0.1% (v/v) formic acid (FA). The eluents were then dried using a SpeedVac™ vacuum concentrator for further immunoaffinity enrichment.

### Immunoaffinity enrichment

A sequential enrichment for each sample was performed using PTMScan® acetyl-lysine (Kac), succinyl-lysine (Ksucc), and propionyl-lysine (Kprop) motif kits (Cell Signaling technology, Inc.) according to the manufacturer’s instruction. Briefly, the dried tryptic peptides obtained above were re-suspended in 1ml of PTMScan® IAP Buffer through votexing and centrifuged at 10,000 *g* for 5 min at 4 °C. Then, the resulting supernatant was used for the first enrichment using Kac motif antibody beads. The peptide solution was mixed with Kac motif antibody beads and incubated for 2 hr on a rotator at 4 °C. After incubation, the beads were collected by centrifugation at 2,000 *g* for 30 s and the supernatant was carefully transferred into a new tube for the second enrichment using Ksucc motif kit. The pelleted beads were sequentially washed twice with cold IAP buffer, three times with H_2_O, and peptides were eluted by adding 55 μl of 0.15% (v/v) trifluoroacetic acid (TFA). The elution was repeated one more time and both eluents were combined for desalting using home-made 10 μm C18 columns (5 mg).

Ksucc and Kprop enrichment was performed using exactly the same procedure as described above for Kac enrichment, except for using the peptide solution following previous enrichment. For Ksucc and Kprop enrichment, the peptide solution was first centrifuged at 2,000 *g* for 30 s to remove any remaining beads from previous enrichment and the supernatant was added to the motif beads directly for enrichment. Similarly, the eluted Ksucc and Kprop peptides were desalted using home-made 10 μm C18 columns (5 mg) and the elutes were dried using a SpeedVac™ vacuum concentrator.

### Mass spectrometry analyses

All enriched peptides were re-suspended in 20 μl 0.1% (v/v) formic acid, and 4 μl of Kac samples and 12 μl of Ksucc and Kprop samples were injected for measurement using high performance liquid chromatography (HPLC) coupled with electrospray ionisation (ESI)-MS/MS. Briefly, peptides were first separated on an reverse phase analytical column (75 μm□×□15□cm; packed with 1.9□μm, 120-Å pore size C18 beads). The mobile phases consisted of 0.1% (v/v) FA in water as buffer A and 0.1% (v/v) FA in acetonitrile as buffer B. A gradient from 5% to 35% buffer B was performed in 120 min at a flow rate of ~300 nL/min. The eluted peptides were then analyzed using Q Exactive HF-X mass spectrometer (ThermoFisher Scientific Inc.) equipped with a nano-electrospray interface operated in positive ion mode. The MS method consisted of one full MS scan from 350 to 1400 m/z followed by data-dependent MS/MS scan of the 16 most intense ions. The full MS was performed in the Orbitrap analyzer with a resolution of 60,000, AGC target of 3e6, and maximum IT of 45 ms, while the MS/MS analyses were performed with a resolution of 15,000, AGC target of 1e5, maximum IT of 50 ms, normalized collision energy (NCE) of 25, and an intensity threshold of 1.6e5. A dynamic exclusion for MS/MS scan was applied with a duration of 20 s. To improve the mass accuracy, all the measurements in Orbitrap mass analyzer were performed with internal recalibration (“Lock Mass”). The charge state rejection function was enabled, with single, unassigned, and ≥4 charge states rejected. All data were recorded with the Thermo Xcalibur™ software (version 3.1) and exported in RAW format.

### Mass spectrometry data processing

Mass spectrometry data processing was performed using MetaLab (https://imetalab.ca/). MetaLab is a versatile and continuously updated software tool for gut microbial protein identification, quantification, as well as downstream metaproteomic data analyses [23, 24]. For this study, a pFind workflow was developed and applied for the identification and quantification of both modified and unmodified peptides/proteins, including lysine acylated ones. Briefly, in the first step, spectra from all raw files were extracted and redundant spectra removed through spectral clustering using PRIDE Cluster [25]. The clustered spectra were then searched against the human fecal microbial Integrated Gene Catalog database (IGC) using Open-pFind (version 3.1.5; http://pfind.ict.ac.cn/) [18]. Open-pFind database search was performed with the following parameters: (1) data extraction with performed with a threshold of −0.5 with mixture spectra option enabled, (2) full cleavage (trypsin KR_C) with up to four missed cleavages was allowed, (3) the tolerance of both the precursor and fragment ions was set to 20 ppm, (4) a peptide length ≥6 amino acids, (5) search mode was set to open search with all modifications in Unimod [26] allowed. The identification results were filtered at peptide level with FDR threshold of 1%. Proteins were kept with ≥1 confidently identified peptides and a FDR <1%. All the resulting protein hits from this first database search were used for generating a refined database. In the second step, the refined database was used for database search for all raw files using Open-pFind with the parameters mentioned above. Quantification of all identified modified or nonmodified peptides was performed using FlashLFQ as described previously. All the identification and quantification results were exported as .txt files for further analysis. Taxonomic assignment and functional annotation of all identified peptides and proteins, respectively, were also carried out and exported as described previously [10, 23].

### Taxonomic analysis

Taxonomic assignment for all identified peptide sequences were performed using a lowest common ancestor (LCA) approach as described previously [23, 27]. For modification-specific taxonomic analysis, only the peptides with the specific modification were used. Relative abundances of taxa at phylum or genus level for each modification data set were calculated using the intensities of distinctive and modified peptides only.

### Functional enrichment

Functional enrichment analysis was performed with hypergeometric probability test using the microbial proteins identified in unenriched samples as background. Briefly, clusters of orthologous group (COG) database was used for functional annotation using DIAMOND (default parameters, e-value = 0.001) [28], and COG id and category were assigned to each identified Kac, Kprop or Ksucc proteins. The numbers of proteins assigned to each COG id or category from modified proteins and background (unmodified proteins) were used for calculating the significance P values of enrichment using hygecdf function in MATLAB (The MathWorks Inc.). P values were corrected using Benjamini-Hochberg adjusted false discovery rate (FDR) using mafdr function in MATLAB. COGs or COG categories with a protein hits more than 10 and FDR-corrected P value less than 0.05 were considered significantly enriched. The enrichment plot was generated using R (version 4.0.2).

### Motif analysis

Motif analysis was performed using pLogo (https://plogo.uconn.edu/) [29]. Briefly, sequence windows with six upstream and downstream amino acids surrounding the modification site were extracted from the database and submitted for pLogo analysis. The total identified microbial proteins from one randomly selected unenriched sample were used as background. Log-odds binomial probability was calculated and plotted with the residues are stacked from most to least represented at each position. Statistical significant threshold (p < 0.05, following Bonferroni correction) was indicated by the red horizontal bar.

### Statistics

Statistical significance of the F/B ratio difference between groups was evaluated using Wilcoxon matched-pairs signed rank test and P < 0.05 was considered significant. Effective of pairing was also evaluated and P < 0.05 was obtained for all comparisons.

## Data and Code Availability

All MS proteomics data that support the findings of this study have been deposited to the ProteomeXchange Consortium via the PRIDE [30] partner repository. The MetaLab 2.1.0 software tool can be downloaded from https://imetalab.ca/. pFind (version 3.1.5) can be downloaded from http://pfind.ict.ac.cn/.

## Acknowledgments

This work was supported by the Government of Canada through Genome Canada and the Ontario Genomics Institute (OGI-156 and OGI-149), the Natural Sciences and Engineering Research Council of Canada (NSERC, grant no. 210034), and the Ontario Ministry of Economic Development and Innovation (ORF-DIG-14405 and project 13440). The authors would like to thank Prof. Si-Min He at the Institute of Computing Technology, CAS, for his support for this project and the comments on the manuscript.

## Supplementary information

**Table S1.** Identified modifications (n ≥ 30) in unenriched and Kac enriched microbiome samples

|  | Modifications | Number of identified peptides |  |
| --- | --- | --- | --- |
|  |  | Kac enrichment | Unenrichment |
| 1 | Acetyl[K] | 57406 | 3 |
| 2 | Carbamidomethyl[C] | 9738 | 11931 |
| 3 | Oxidation[M] | 6173 | 2882 |
| 4 | Deamidated[N] | 2240 | 3018 |
| 5 | Propionyl[K](Delta_H(4)C(3)O(1)[K]) | 1263 | 0 |
| 6 | Gln->pyro-Glu[AnyN-termQ] | 338 | 1 |
| 7 | Cation_Fe[II][D] | 276 | 294 |
| 8 | Cation_Fe[II][E] | 164 | 0 |
| 9 | Formyl[K] | 155 | 5 |
| 10 | Carbamyl[K] | 79 | 0 |
| 11 | Dioxidation[W] | 46 | 0 |
| 12 | Pyro-carbamidomethyl[AnyN-termC] | 43 | 11 |
| 13 | Gly->Pro[G] | 36 | 0 |
| 14 | Val->Xle[V] | 36 | 0 |
| 15 | Butyryl[K](Crotonaldehyde[K]) | 31 | 1 |
| 16 | Asp->Asn[D] | 30 | 0 |
| 17 | Glu->Asp[E] | 19 | 124 |
| 18 | Xle->Val[I] | 14 | 370 |
| 19 | Methyl[D](Asp->Glu[D]) | 14 | 71 |
| 20 | Thr->Ala[T] | 14 | 63 |
| 21 | Cation_Ca[II][E] | 12 | 30 |
| 22 | Ala->Thr[A] | 9 | 60 |
| 23 | MesitylOxide[K] | 9 | 42 |
| 24 | Ala->Ser[A] | 8 | 400 |
| 25 | Cation_Ca[II][D] | 7 | 52 |
| 26 | Deoxy[S](Ser->Ala[S]) | 6 | 41 |
| 27 | Acetyl[AnyN-term] | 4 | 35 |
| 28 | Cation_Na[D] | 3 | 51 |
| 29 | Ala->Val[A] | 3 | 39 |
| 30 | Methyl[S](Ser->Thr[S]) | 2 | 135 |
| 31 | Glu->pyro-Glu[AnyN-termE] | 2 | 55 |
| 32 | Ammonia-loss[N] | 2 | 35 |
| 33 | Carboxymethyl[K] | 1 | 60 |
| 34 | Delta_H(6)C(6)O(1)[K] | 1 | 48 |
| 35 | Glu->Gln[E] | 0 | 69 |
| 36 | Carboxyethyl[K] | 0 | 37 |
| 37 | Xlink_BS2G[113][K] | 0 | 32 |

**Table S2** All significantly enriched COGs by lysine acylated proteins in microbiome Table S2.xlsx

## Supplementary Figures

**Figure S1.**
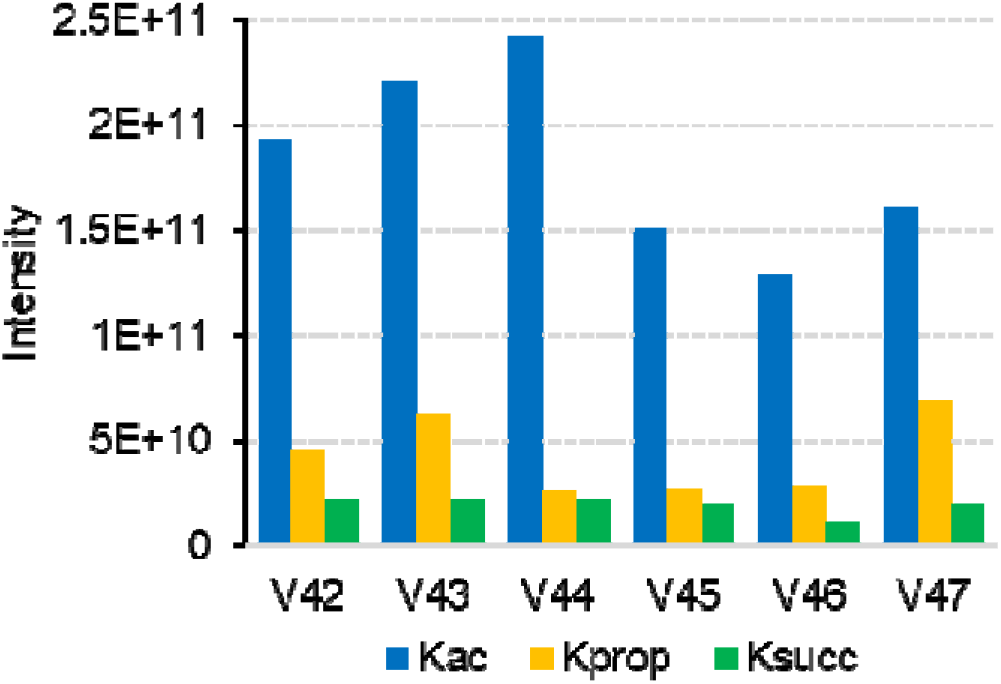
Estimation of the abundance of lysine acylations in human gut microbiome. The sum intensity of all identified Kac, Kprop or Ksucc peptides was used as a proxy for the abundance of Kac, Kprop and Ksucc, respectively.

**Figure S2.**
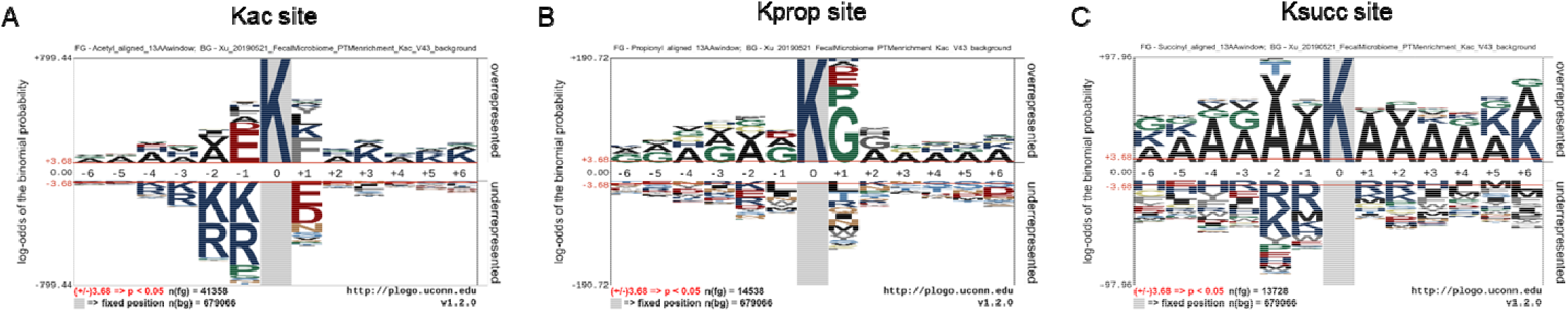
Motif of identified lysine acetylation, propionylation and succinylation sites. Motif analysis was performed using pLogo (https://plogo.uconn.edu/) with figures directly exported from the website. Log-odds binomial probability was plotted with the residues being stacked from most to least represented at each position. Significance threshold (p < 0.05, following Bonferroni correction) was indicated by the red horizontal bar.

**Figure S3.**
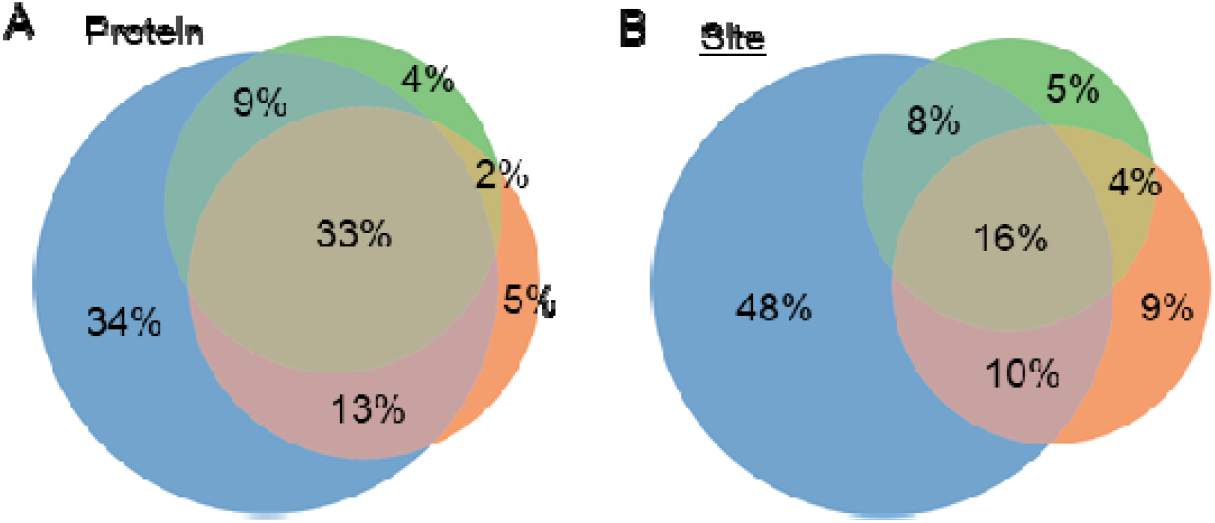
Overlap of identified lysine acylated proteins and site.

**Figure S4.**
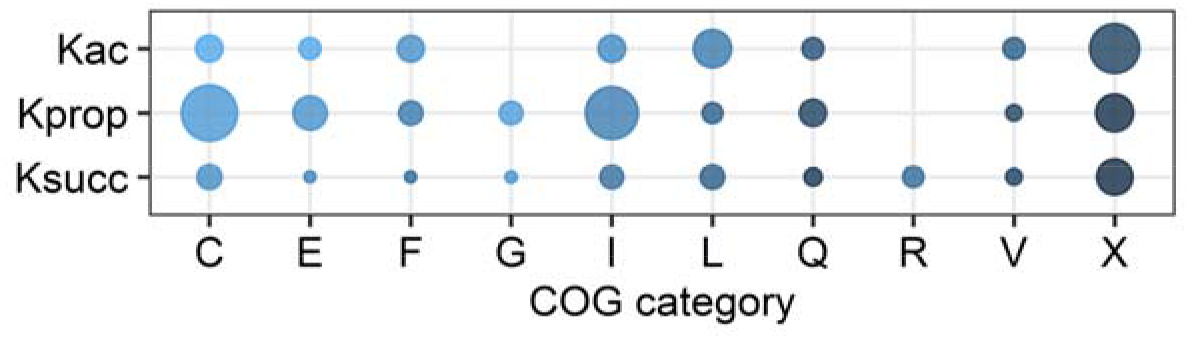
Significantly enriched COG categories by Kac, Kprop or Ksucc proteins identified in microbiome. The size of the dot indicates the –log10(P value) and color indicates the number of hits (low, light blue; high, dark blue). C, Energy production and conversion; E, Amino acid metabolis and transport; F, Nucleotide metabolism and transport; G, Carbohydrate metabolism and transport; I, Lipid transport and metabolism; L, Replication and repair; Q, Secondary metabolites biosynthesis, transport and catabolism; R, General function prediction only; V, Defense mechanisms; X, mobilome.

